# New mitochondrial gene order arrangements and evolutionary implications in the subclass Octocorallia

**DOI:** 10.1101/2024.06.15.599068

**Authors:** Angelo Poliseno, Andrea M. Quattrini, Yee Wah Lau, Stacy Pirro, James D. Reimer, Catherine S. McFadden

## Abstract

The complete mitochondrial genomes of octocorals typically range from 18.5 kb to 20.5 kb in length, and include 14 protein coding genes (PCGs), two ribosomal RNA genes and one tRNA. To date seven different gene orders (A-G) have been described, yet comprehensive investigations of the actual number of arrangements, as well as comparative analyses and evolutionary reconstructions of mitochondrial genome evolution within the whole subclass Octocorallia have been often overlooked. Here we considered the complete mitochondrial genomes available for octocorals and explored their structure and gene order variability. Our results updated the actual number of mitochondrial gene order arrangements so far known for octocorals from seven to twelve, and allowed us to explore and preliminarily discuss the role of some of the structural and functional factors in the mitogenomes. We performed comparative mitogenomic analyses on the existing and novel octocoral gene orders, considering different mitogenomic structural features such as genome size, GC percentage, AT- and GC-skewness. The mitochondrial gene order history mapped on a recently published nuclear loci phylogeny showed that the most common rearrangement events in octocorals are inversions, and that the mitochondrial genome evolution in the subclass is discontinuous, with rearranged gene orders restricted only to some regions of the tree. We believe that different rearrangement events arose independently and most likely that new gene orders, instead of being derived from other rearranged orders, came from the ancestral and most common gene order. Finally, our data demonstrate how the study of mitochondrial gene orders can be used to explore the evolution of octocorals and in some cases can be used to assess the phylogenetic placement of certain taxa.

## Introduction

The recovery of complete mitochondrial genome sequences for taxonomic studies and for biogeographic, phylogenetic and evolutionary investigations has become routine over the last decade (Plazzi et al. 2016; Tan et al. 2019; Reyes-Velasco et al. 2021). The increase in available complete mitogenomes has been driven by an overall lowering of the sequencing costs (Nunez and Oleksiak 2016), by the effectiveness of the molecular technologies employed, and by availability of bioinformatic tools for mitogenome analyses (Forni et al. 2019; Hoban et al. 2022). Despite the mitogenomic approach becoming popular in evolutionary biology, some taxa are still poorly sampled. For example, within the subclass Octocorallia, the number of complete mitogenome sequences has only recently increased, with about 65% of the total sequences publicly available being deposited within the last four years. To date seven different gene order arrangements have been reported within the subclass (Brugler and France 2008; Uda et al. 2011; Brockman and McFadden 2012; Pante et al. 2013), yet, as recently shown for pennatulaceans (Hogan et al. 2019), new rearrangements are still being discovered in different species.

The first studies dealing with mitochondrial gene arrangements in octocorals identified the presence within the mitogenome of four conserved gene blocks whose inversion or translocation could have potentially led to five of the observed rearrangements (A-E) (Uda et al. 2011; Brockman and McFadden 2012; Figueroa and Baco 2015). The occurrence of octocoral mitogenome rearrangements that do not conserve these four gene blocks was preliminarily proposed by Pante et al. (2013) after the screening of gene junctions in a calcaxonian species (*Isidoides armata*; arrangement F). Later, Shimpi et al. (2017) showed that transcriptional units encompass genes from different gene blocks, and more recently Hogan et al. (2019) found a non-conserved gene block within the mitogenome of an undescribed *Umbellula* species (arrangement G). Recently, unique novelties in the mitogenome structure of octocorals were discovered, including a bipartite genome within pennatulaceans (Hogan et al. 2019) and the lack of a *mtMutS* gene in the mitogenome of *Pseudoanthomastus* sp.1 (Muthye et al. 2022). Despite such recent efforts, a comprehensive overview of the different gene order arrangements and estimations of genome arrangement history to date have been poorly explored. Regarding the mitochondrial gene order arrangements so far described, Brockman and McFadden (2012) hypothesized that the most common arrangement (A) can be considered ancestral among octocorals due to its broad phylogenetic distribution within the class Octocorallia. However, ancestral state reconstruction analysis has never been done to confirm this hypothesis.

Size, structure and organization of mitochondrial genomes aside, mitochondrial protein-coding genes (PCGs) are frequently used for phylogenetic inference and for biogeographic and historical evolutionary reconstructions (Gissi et al. 2008; Osigus et al. 2013). This is also true for octocorals which, in spite of having a highly reduced mitochondrial mutation rate (Shearer et al. 2002; Bilewitch and Degnan 2011), have had their mitogenome sequences exploited widely in phylogenetic and evolutionary studies (Kayal et al. 2013; Poliseno et al. 2017; 2017a; 2021; Ramos et al. 2023; Feng et al. 2023).

In this study, we reconsidered the complete mitogenomes available on GenBank and added the sequences of three *Phenganax* species (*P. marumi, P. subtilis* and *P. stokvisi*) for comparative and evolutionary analyses. We mapped the mitochondrial gene arrangement history on a recently published phylogeny and reported on ancestral mt-genome states using nine of 12 available mitochondrial genome arrangements.

## Material and methods

### Specimens, DNA extraction, library preparation and DNA sequencing

The majority of the specimens used for this study were previously sequenced (Quattrini et al. 2018, 2020) and their complete mitogenome sequences are available on GenBank (OL616196 – OL616289). Mitogenomes for three species of the genus *Phenganax* (family Acrossotidae) were newly sequenced in this study. DNA extraction of *Phenganax* spp. was performed using a Qiagen DNeasy Blood & Tissue kit (Qiagen, Tokyo). DNA extracts were sent to Iridian Genomes (Bethesda, MD, USA) for library preparation and for genome skimming using the Illumina HiSeq X Ten sequencer.

The raw reads for *Phenganax* spp. (SRR12278765, SRR12621204 and SRR12626619) were trimmed and filtered with Trimmomatic 0.39 (Bolger et al. 2014). Mitogenome assembly was performed with default parameters in NOVOPlasty 4.3.1 (Dierckxsens et al. 2016) using the partial *mtMutS* sequence of *P. subtilis* (MN164586) as a starting sequence. Mitogenome annotation was performed with the software Geneious prime^®^ 2022.02 (Kearse et al. 2012) using the ORFinder function and the complete sequence of other Acrossotidae as a reference. The sequences obtained were deposited on GenBank under accession numbers PP330783-PP330785.

In order to confirm the novel gene arrangements in *Keroeides* (OL616243) and *Muricella* (OL616247) suggested by the mitogenome reconstructions of Muthye et al. (2021), we designed PCR primers to amplify and sequence regions of the mitogenome that span gene junctions that are not found in the canonical arrangement (A). Primers were designed to amplify across junctions between *Cox3*-*Nad3, Cob*-*Nad4L, Nad3*-*Atp6* and *Nad4*-*Cox3* in *Keroeides*, and across *Cox2*-*Nad4, Cox2*-*Nad5, Cox3*-*Nad4* and *Cox3*-*Cob* in *Muricella*. Following successful PCR, amplicons were Sanger sequenced to confirm the sequence matched the annotated mitogenome.

### Comparative analyses, mitochondrial gene order arrangements and gene order history reconstruction

For comparative analyses, the base composition, GC percentage, and AT/GC skew were calculated with PhyloSuite 1.2.3 (Zhang et al., 2020). AT- and GC skewness were calculated using the equations: AT-skew = (A-T)/(A+T) and GC skew = (G-C)/(G+C) (Perna and Kocher 1995). In summary, AT- and GC-skewness may represent differences between two strands due to asymmetries in the mitogenome replication process, in which one strand ‘prefers’ C/A over G/T.

Analyses of mitogenomic rearrangements based on common intervals were conducted with CREx (Bernt et al. 2007) using default parameters. CREx uses a data structure called strong common interval tree (Bérard et al. 2007) and, with the help of a distance matrix and interval tree, heuristically determines genome rearrangement scenarios between the given gene orders. Ancestral state reconstruction of mitochondrial gene order was performed using a parsimony ancestral state method with default parameters in Mesquite 3.81 (Maddison and Maddison 2023) considering the phylogenetic tree inferred from nuclear loci by Quattrini et al. (2023) as the input file. The starting tree was slightly modified by removing *Corymbophyton, Leptophyton* and *Tenerodus*─ which are part of a larger dataset of unpublished data─ and it was transformed into a binary ultrametric tree. The use of the nuclear loci tree for character-mapping and ancestral state reconstructions was preferred over mitochondrial tree reconstructions as evidenced by the recent study of Ramos et al. (2023) that pointed out the impact of selection on the evolution of octocoral mitogenomes, and by Quattrini et al. (2023) who highlighted evident limitations when inferring accurate species relationships using complete mitochondrial genomes likely due to the lack of neutral evolution, rate variability and rapid introgression. In addition, the different gene order arrangements were mapped on the same nuclear phylogeny with all the allowed consistency methods (i.e. strong, weak and parsimonious weak) in TreeREx (Bernt et al. 2008).

## Results

### Mitochondrial genome structure and gene order arrangements

For *Keroeides* (gene order J) we successfully amplified sequences spanning three gene blocks, including *Cob*-*Nad4L, Nad4*-*trnM*-*Cox3* and *Nad6*-*Atp6*, but we were unable to validate the juxtaposition of *Cox3-Nad3* (see Fig.1). We were only able to verify a portion of the new gene order proposed for *Muricella* (gene order K), as multiple attempts to sequence across the *Nad4-Cox2* junction using different primer pairs failed. Moreover, a new genome assembly from genome-skimming data (Quattrini et al. 2024) juxtaposed *Cox3* and *Nad4*, a junction that was subsequently verified using PCR. These results suggest the *Muricella* genome assembly published by Muthye et al. (2022) is incorrect.

In addition to the seven gene order arrangements already described for octocorals (A-G), we detected five new arrangements, which have been here designated as follows: ‘F1’(*Acrossota amboinensis* – OL 616200; *Arula petunia* – OL616211; *Paratelesto* sp. – OL16258; *Tubipora* sp. – OL616283 and *Phenganax stokvisi* – PP330784), ‘H’ (*Phenganax subtilis* – PP330785 and *P. marumi* – PP330783), ‘I’ (*Anthelia glauca* – OL616207; *Caementabunda simplex* – Ol616216, *Coelogorgia palmosa* – OL6162224; *Ovabunda macrospiculata* – OL616252 and *Protodendron repens* – OL616263), ‘J’ (*Keroeides* – OL6162439) and ‘K’ (*Muricella* sp. – OL61624) (see Figure 1). In terms of gene content, the new arrangements have the same number of PCGs, ribosomal RNA and tRNA genes as the mitogenomes described so far. The mitogenome sizes were also consistent with those of other octocorals and typically ranged from 18Kb to 20Kb.

**Figure 1.**
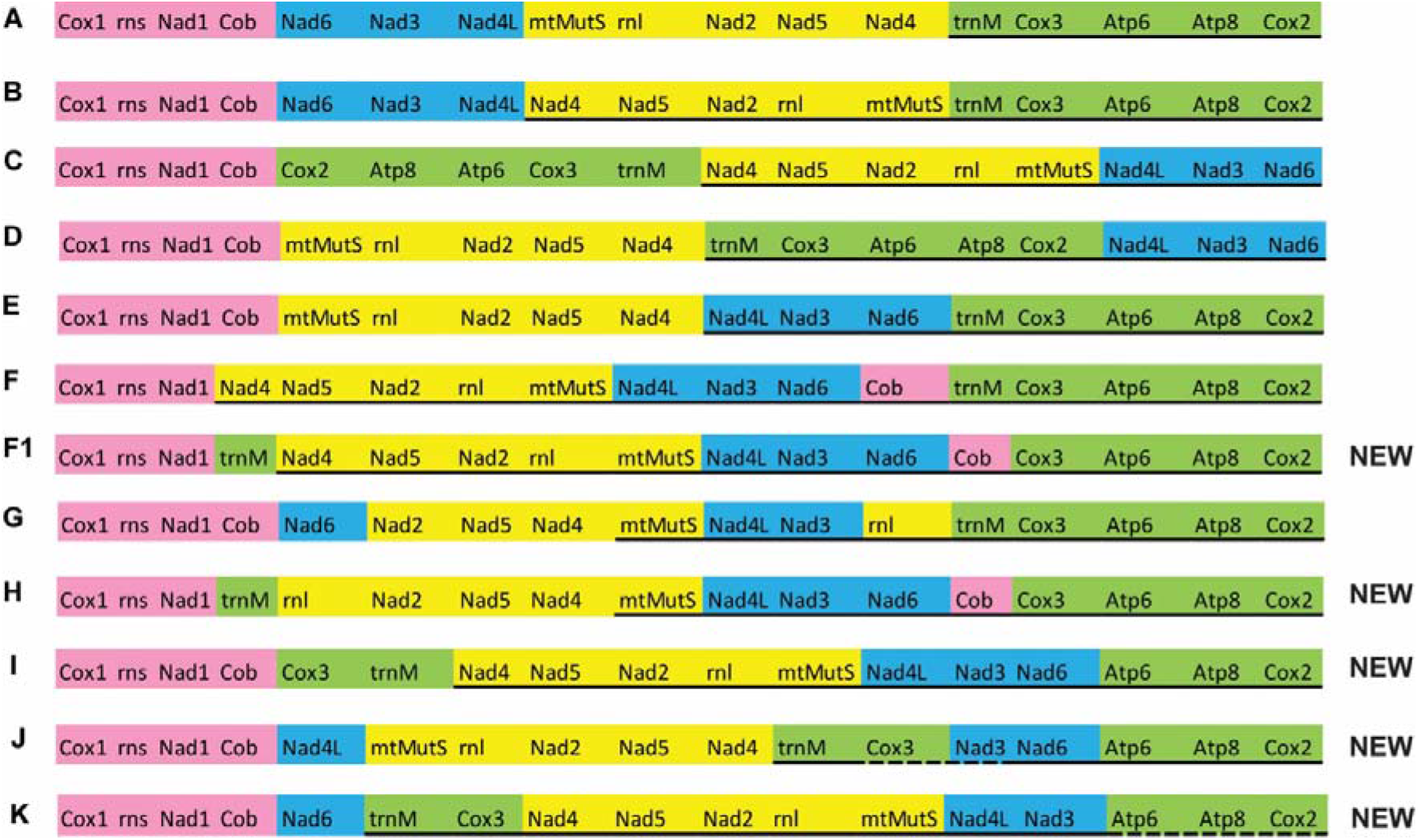
Summary of the different mitochondrial gene arrangements discovered for octocorals. Conserved blocks of genes as for Brockman and McFadden (2012) are marked with diverse colours. Letters on the left refer to gene arrangements following previous schemes. Solid lines below gene names indicate genes encoded on light strand. The dashed line below gene names in the arrangements ‘J’ and ‘K’ indicates that the placement of these genes in the mitogenome was not corroborated by Sanger sequencing.

The majority of the specimens investigated had gene order ‘A’, which was shared across the phylogenetic tree among most families of orders Malacalcyonacea and Scleralcyonacea. The gene order arrangement ‘B’ was shared by some members of the family Keratoisididae and by a single genus (*Anthoptilum*) within family Anthoptilidae. Gene order arrangements ‘C’ and ‘D’ were typical among the coralliid genera *Anthomastus, Hemicorallium, Paragorgia* and *Pleurocorallium*. Gene order arrangements ‘F1’and ‘H’ have been found among *Acrossota, Arula, Paratelesto, Phenganax* and *Tubipora* species, whereas type ‘I’ was restricted to the family Xeniidae and its sister taxon *Coelogorgia palmosa*. The remaining arrangements currently only involve single species, for instance type ‘E’ (*Paraminabea aldersladei*), type ‘F’ (*Isidoides armata*), type ‘G’ (*Umbellula* sp. 1), type ‘J’ (*Keroeides* sp.) and type ‘K’ (*Muricella* sp.). All the newly discovered arrangements have undergone the inversion of PCGs across different gene blocks. In particular, compared to the most common gene order, ‘A’, we found that ‘F1’ and ‘I’ have undergone inversion of ten PCGs across four and three gene blocks, respectively. Within type ‘F1’ trnM has been inverted and placed between *Nad1* and *Nad4*, whereas *Cob* had *Nad6* and *Cox3* as flanking genes. Regarding type ‘I’ the block including *trnM, Cox3, Atp6, Atp8* and *Cox2* has seen the separation of *Cox3* and *trnM* from the other genes of the block and their inversion between *Cob* and *Nad4*. Type ‘J’ had two PCGs inverted in one gene block, with *Nad4L* being placed between *Cob* and *mtMutS*, and the other two PCGs of the block inverted within the *trnM*-*Cox3*-*Atp6*-*Atp8*-*Cox2* block. In arrangement ‘H’, *trnM* and *Cob* have been inverted and moved between *Nad1* and *16S rRNA* and between *Nad6* and *Cox3*, respectively. Concerning arrangement ‘K’, *Nad6* is flanking *Cob* and *Cox3*, whereas the five PCGs within the *mtMutS*-*Nad4* block were inverted. (Figure 2).

**Figure 2.**
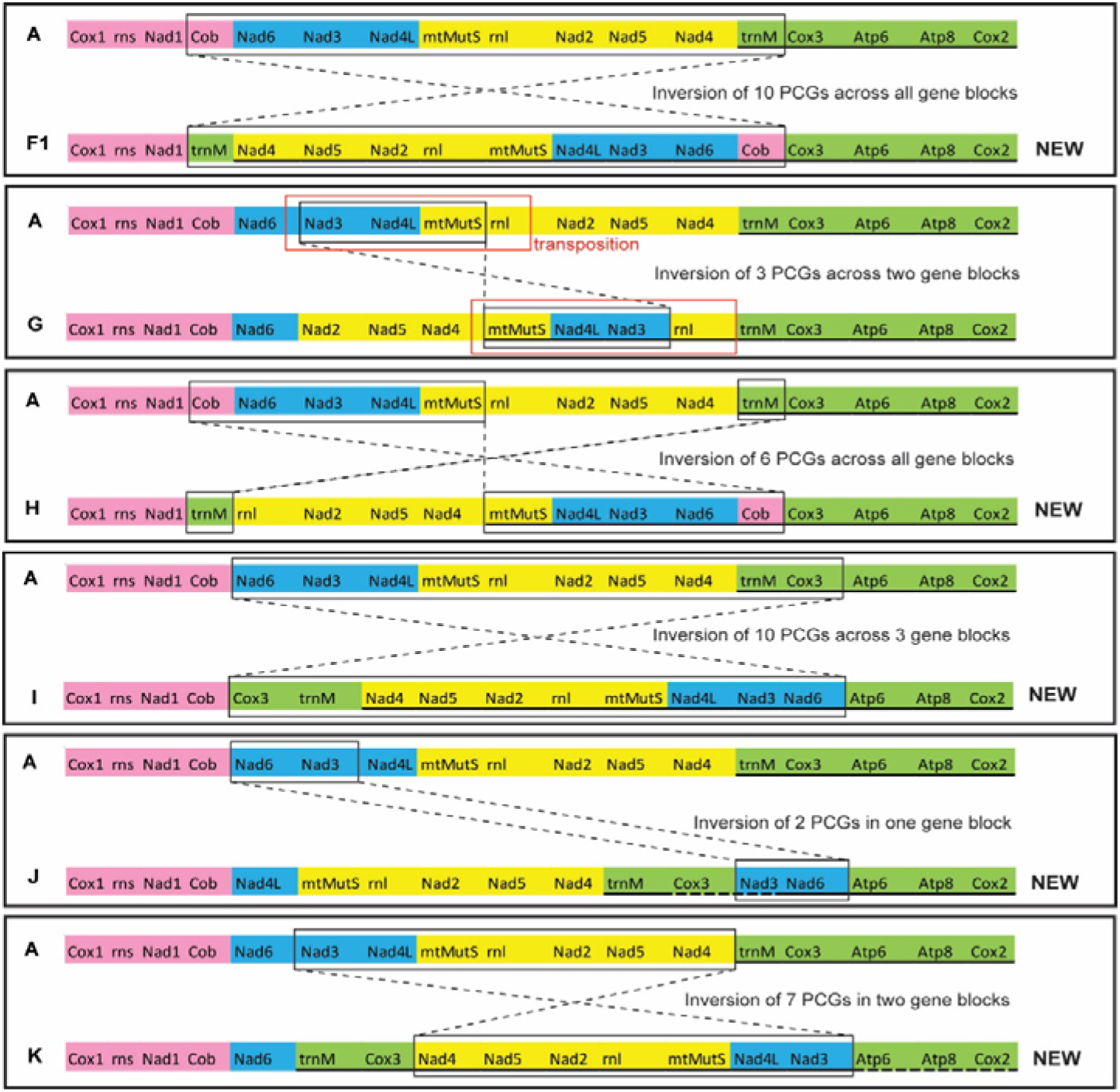
Scheme of changes (inversions and transpositions) that occurred in the different gene order arrangements. Conserved blocks of genes as for Brockman and McFadden (2012) are marked with diverse colours. Letters on the left refer to gene arrangements following previous schemes. Solid lines below gene names indicate genes encoded on light strand. The dash line below gene names in the arrangements ‘J’ and ‘K’ indicates that the placement of these genes in the mitogenome was not corroborated by Sanger sequencing.

Among the octocorals investigated, mitogenome sizes ranged from 18,052 bp (*C. pabloi*) to 20,246 bp (*C. granulosa*). Most of the octocoral species with a gene order different from ‘A’ had an average genome size (18.5 Kb – 19.5 Kb), yet four of the seven species with a larger genome─ due to longer intergenic regions─ had atypical gene order arrangements (e.g. *P. aldersladei* (E), *A. amboinensis* and *Paratelesto* sp. 1 (F1) and *P. subtilis* (H)) (Figure 3A). Concerning GC content, octocorals have percentages ranging from 35.5% to 38.5%. This range is shared by most of the species analysed here, with the exception of a group of specimens having a GC content greater than 38.5% and 11 species with a GC content lower than 35.5% (Figure 3B, Supplementary Table S1).

**Figure 3.**
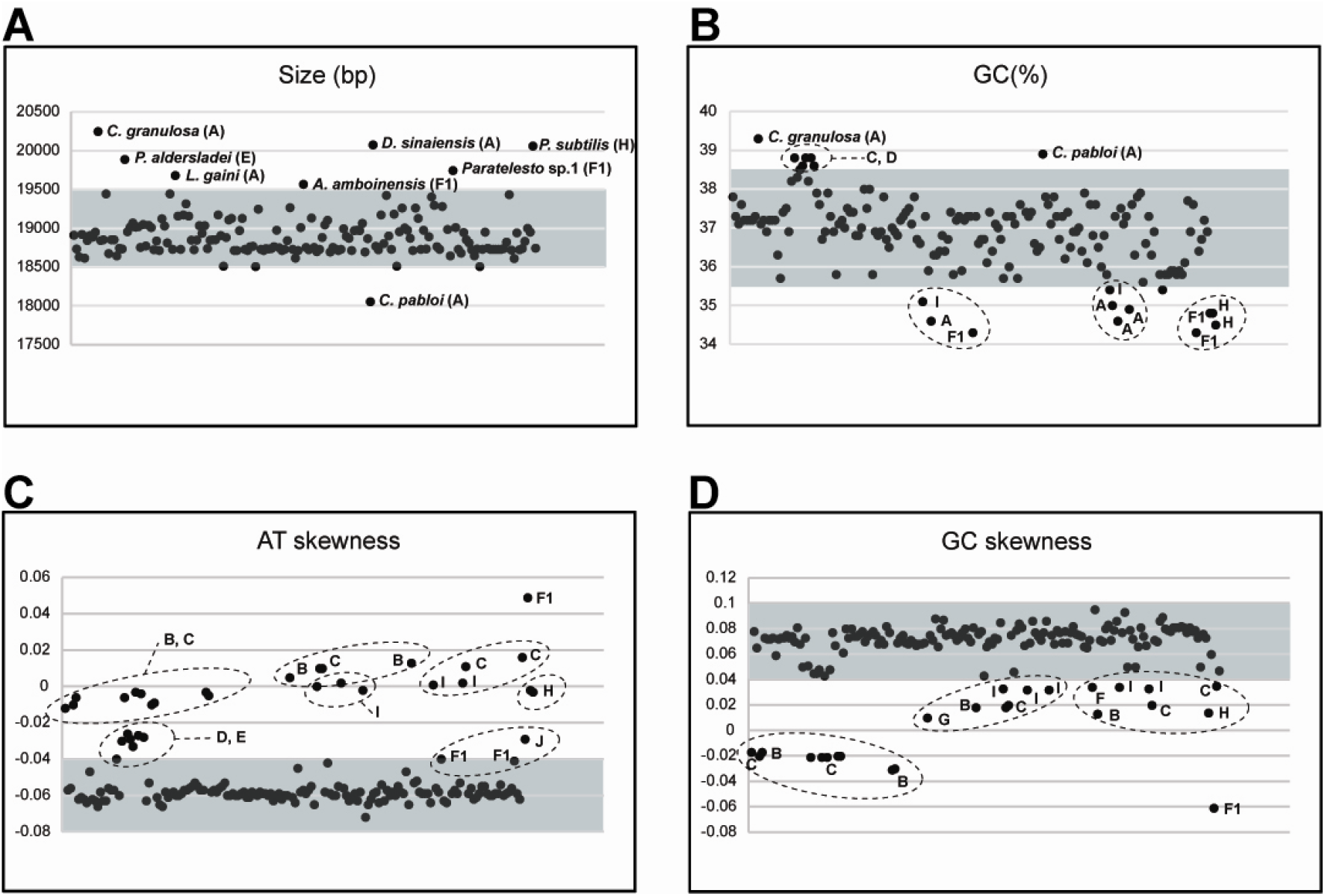
Statistics on the mitochondrial genomes of 174 octocorals including genome size (A), GC percentage (B), AT-skewness (C) and GC skewness (D). Letters from A to J refer to the different gene order arrangements. Grey areas in each graph indicate the most common range of values for the dataset.

In agreement with Feng et al. (2023), who explored the AT- and GC skewness among cnidarians, we found that the majority of the specimens investigated had negative AT skewness, comprising values between -0.08 and -0.04. Interestingly, all specimens with values falling out of the range (> -0.04) had gene order arrangements other than A (Figure 3C). Among the specimens with either positive AT skewness or with values close to zero, the most common gene order arrangements were B, C, F1 and I. *Phenganax stokvisi* (F1) had the greatest AT skewness value (0.049) (Figure 3C). Most octocoral species analysed had positive GC-skewness and, similar to what was found for AT skewness, all specimens with GC skewness values out of the normal range (0.04-0.1) had uncommon gene order arrangements such as B, C, F, F1, G, H and I (Figure 3D). These data indicate that octocoral mitogenomes are biased toward using GC rather than AT.

### Mitogenome rearrangement mapping on nuclear loci phylogeny

Overall our TreeRex analysis inferred 11 inversion events, one transposition and one inverse transposition, of which six occurred within the order Scleralcyonacea and seven within the order Malacalcyonacea (Figure 4). All the rearrangement events occurred at the shallower nodes of the tree. Among the scleralcyonaceans investigated, clades S1 and S3 included taxa with a single gene order (A), whereas among malacalcyonaceans five of eight clades comprised members with gene order A only. Even though our phylogenetic analyses did not include representatives for each of the twelve mitogenome arrangements (arrangements D, G, H were not included) we were able to gain information about the different mitogenome(s) structure and gene orders across the subclass. For instance, the taxonomic group with the highest proportion of rearrangements and therefore with the greatest number of gene arrangements was the family Coralliidae, including four gene orders (A; C; D; E).

**Figure 4.**
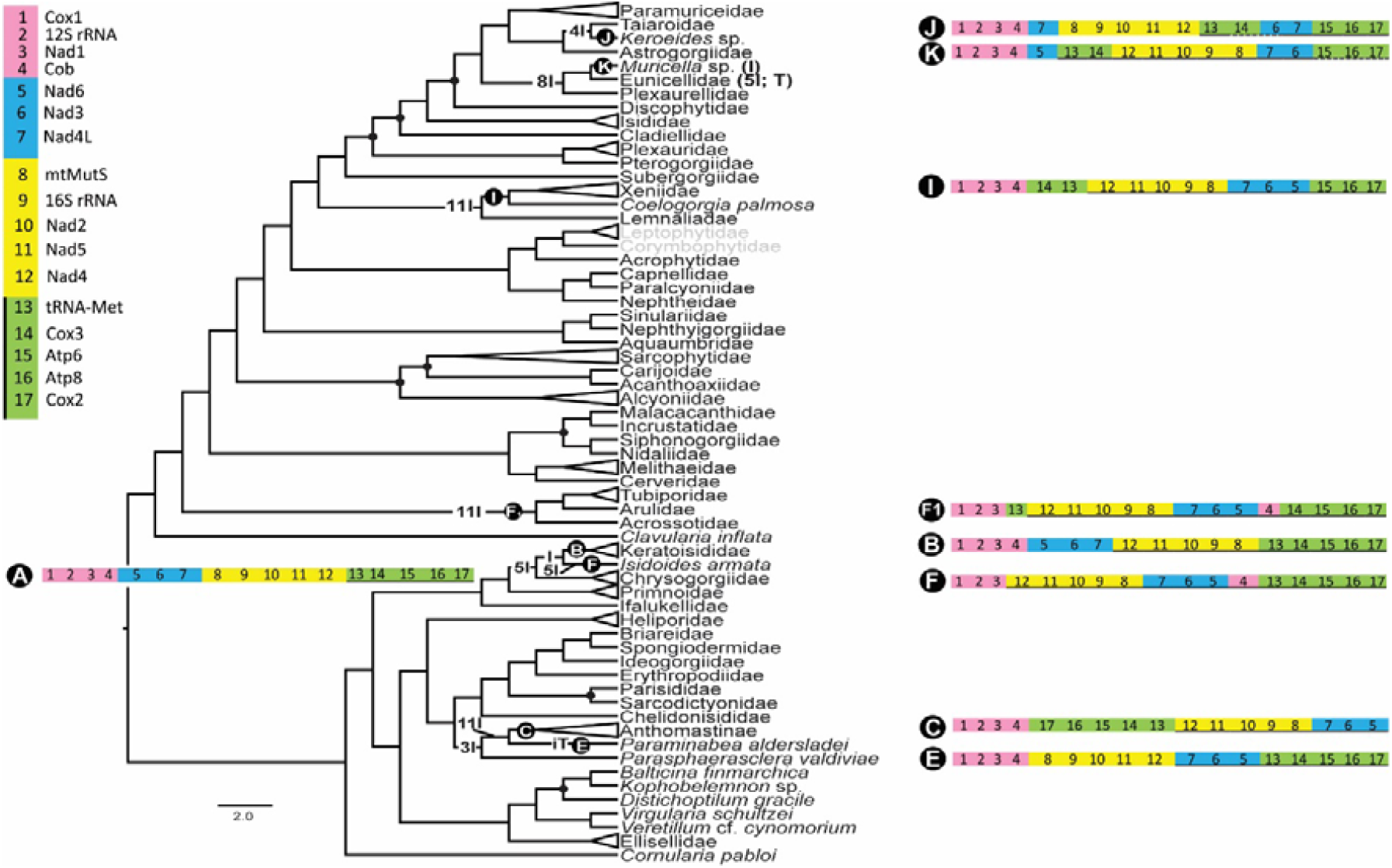
Gene order evolution of the subclass Octocorallia based on TreeRex analyses mapped on the nuclear loci tree by Quattrini et al. (2023). In order to facilitate readability of the tree, many clades were reduced to family level. Gene orders are shown on the different nodes of the tree and the most likely evolutionary scenario(s) leading to the new arrangements are provided either at the different nodes or next to the species names. Members of families Leptophytidae and Corymbophytidae (in grey) were not considered for the gene order scenario as they are part of a larger dataset to be analysed in a different publication. I = Inversion; T = Transposition; iT = inverse Transposition.

Our ancestral state reconstruction supported gene order ‘A’ as ancestral within Octocorallia (Fig. 5). Among scleralcyonaceans, the common ancestral mitogenome arrangement of Chrysogorgiidae, Keratoisididae and *Isidoides armata* underwent three separate inversion events leading to arrangement types B and type F, respectively (Figure 4, Figure 5). The common ancestral mitochondrial gene order of *Parasphaerasclera valdiviae*, Anthomastinae and *Paraminabea aldersladei* underwent two separate events; one leading to type C in Anthomastinae and the second leading to type E in *Paraminabea aldersladei*. Regarding malacalcyonaceans, the common ancestral mitogenome arrangement of Acrossotidae, Arulidae and Tubiporidae underwent eleven inversions in a single event leading, for all families, to arrangement F1 (Figure 4, Figure 5). The common ancestral gene order type I of Xeniidae and *Coelogorgia palmosa* was derived from type A as a single inversion rearrangement event (Figure 4). The common ancestral gene order of Astrogorgiidae, Taiaroidae and *Keroeides* sp. underwent a single rearrangement event consisting of four inversions that led to arrangement type J in *Keroeides* sp.

**Figure 5.**
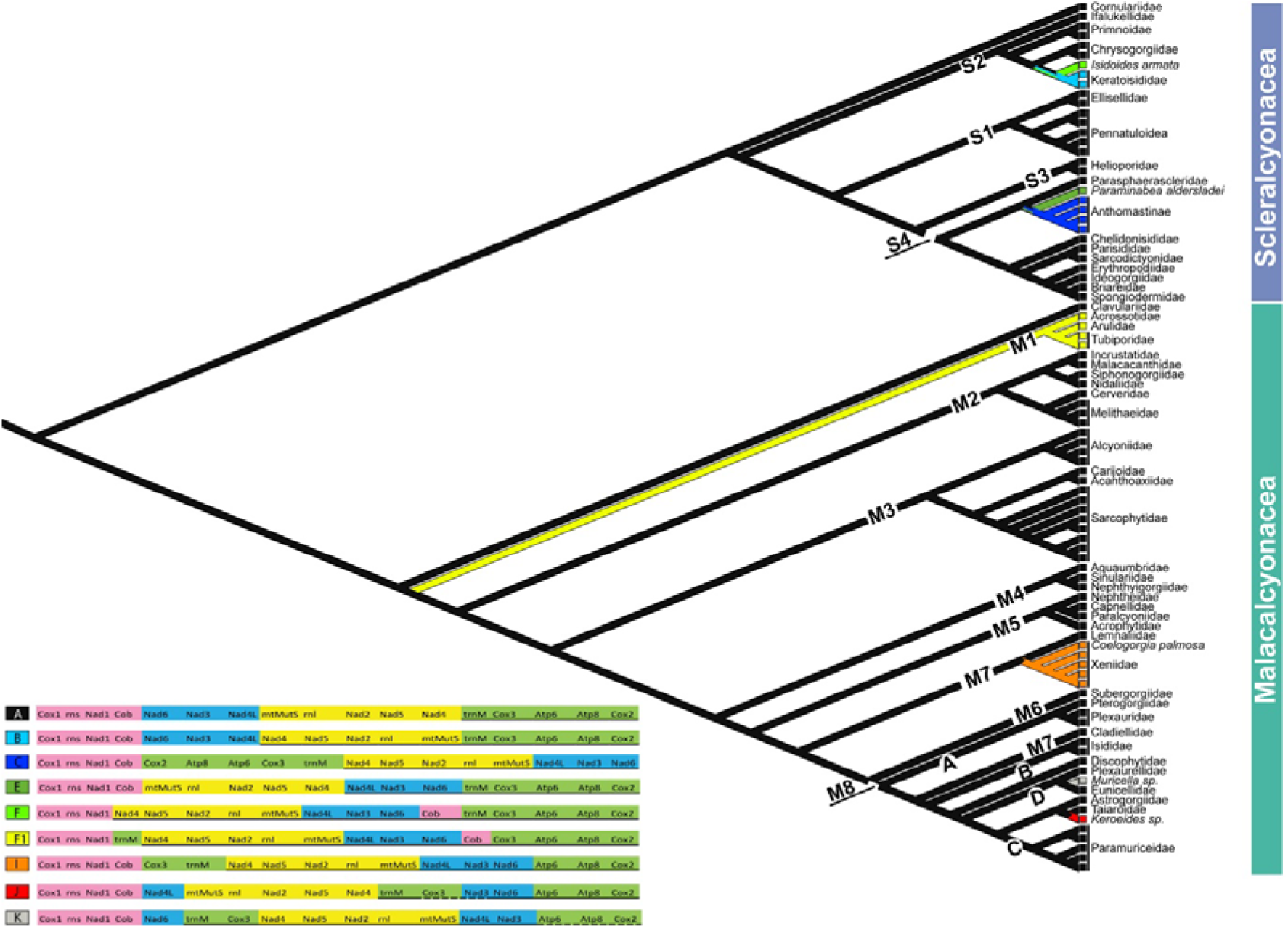
Ancestral state reconstruction of nine of the 12 mitochondrial gene order arrangements of the subclass Octocorallia. In order to facilitate readability of the tree, many clades were reduced to family level. Division into four Scleralcyonacean groups (S1-S4) and eight Malacalcyonacean groups (M1-M8) follows the systematic scheme proposed by McFadden et al. (2022).

Interestingly, among the taxa investigated, the only genera reporting intrageneric variation at the level of gene order were *Umbellula* (*U*. sp. 1 – type G; *U*. sp. 2 – type A) and *Phenganax* (*P. stokvisi* – type F1; *P. marumi* and *P. subtilis* – type H). However, we expect similar results in other groups as additional mitogenomes within species-rich genera are sequenced.

## Discussion

Our overview on the mito-gene order arrangements of the subclass Octocorallia showed that the actual number of known mitogenome arrangements is 12, and that approximately 21% of species investigated had a different gene order than the common ancestral one (A), which is shared across the families of both orders. Our data revealed that except for families Acrossotidae, Arulidae and Tubiporidae (clade M1), Coelogorgiidae and Xeniidae (clade M7), and Anthogorgiidae and Keroeididae (clade M8), the remaining malacalcyonacean families have the ancestral gene order arrangement, and that five (A, F1, H, I and J) of the 12 arrangements are found among members of this order. Seven arrangements (A-G) have been detected among scleralcyonaceans, with the ancestral one (A) being the only type shared by species of both orders. We observed that atypical gene order arrangements within Scleralcyonacea are restricted to a few groups such as sea pens (*Umbellula* and *Anthoptilum*), bamboo corals (*Isidoides* and Keratoisididae), and coralliids (e.g., *Anthomastus, Corallium, Hemicorallium, Paragorgia*; *Paraminabea* and *Pleurocorallium*) and, except for *Paraminabea*, all of these genera are found in deep waters. Our data on scleralcyonaceans suggest that mitogenome rearrangements frequently occur in deep-sea species, as has already been proposed for other metazoans such as Annelida and Holothuria (Zhang et al. 2018; Tempestini et al. 2020; Sun et al. 2021). However, most scleralcyonaceans inhabit deep waters, so the proportion of deepwater species with gene arrangements likely reflects where most members of that order can be found. Correlation between species’ depths and mitogenome re-arrangements was only partly supported by the data obtained on malacalcyonaceans. In fact, *Kereoides* is the only genus with an atypical mitochondrial arrangement (type J) inhabiting deep-sea waters, whereas both xeniids (type I) and members of Acrossotidae, Arulidae and Tubiporidae (types F1 and H) are exclusively found in shallow waters. However, not all genera and species inhabiting deep waters have mitogenome rearrangements, as shown for example by chrysogorgiids and primnoids, which have instead the ancestral arrangement. Unlike octocorals, the mitogenomes of other anthozoan orders such as Antipatharia, Scleractinia and Zoantharia have highly conserved gene orders and content. In particular, among scleractinians, the majority of the mitogenomes so far sequenced have the same gene order, with rearrangements reported only in five genera, of which three (*Desmophyllum, Madrepora, Solenosmilia*) occur in deep waters (Seiblitz et al. 2022). Despite Antipatharia and Zoantharia including deep-water species, these orders have extremely low levels of mitogenome rearrangements (Antipatharia; Barrett et al. 2020) or no rearrangements at all (Zoantharia; Poliseno et al. 2020).

Living in deep environments seems inappropriate to explain these octocoral mitochondrial rearrangements, and thus other factors should be taken into account. Some of these rearrangements may have biological causes, as suggested for bilaterians (Bernt et al. 2013), while for non-bilaterians, substitution rates, oxidative stress, the presence of tRNAs, and increasing mitogenome sizes are recognized as some of the principal factors that can contribute to mitogenome rearrangements (Saccone et al. 1999; Luo et al. 2015; Tempestini et al. 2018; Zhang et al. 2018; Sun et al. 2021). However, octocoral mitogenomes are known to have slow nucleotide substitution rates (Muthye et al. 2022) due to the presence of the putative DNA mismatch repair function of *mtMutS* (Bilewitch and Degnan 2011), and thus the increase of substitution rates to explain the overall great variety of mitogenome rearrangements does not seem to be the case for these organisms, but can possibly explain the placement of some of these groups on long branches in the phylogenetic trees. Regarding oxidative stress, mtDNA damage may lead to different major changes within the mitogenome that can be converted either to point mutations or to mitogenomic rearrangements (Kajander et al. 2000; Dowton and Campbell 2001). Octocoral *mtMutS* seems capable of counteracting the effects of oxidative stress (Shimpi et al. 2017), drastically reducing the occurrence of mitogenome rearrangements. Transfer RNAs have been thought to act as mobile elements enabling rearrangements (Dowton and Campbell 2001; Luo et al. 2015), but the paucity of tRNAs in octocoral mitogenomes ─ tRNA^Met^ is the only tRNA detected for all octocorals─ cannot alone justify the high number of rearrangements discovered within the subclass. Recent studies on Annelida (Sun et al. 2021; Struck et al. 2023) suggest that the increase of mitogenome size due to the presence of extended intergenic regions (IGRs) and large sequence duplication was not necessarily correlated to new mitogenome rearrangements. Similarly, our data suggested that only four species with mitogenome rearrangements presented a genome size larger than the typical octocoral average. While the above factors do not appear to be correlated with the onset of new gene orders in octocorals, our data regarding GC percentages and AT- and GC-skewness highlighted interesting patterns. The majority of the species with a GC percentage, AT- and GC-skewness over or above the octocoral averages have a gene order other than the ancestral one (A). Therefore, our findings show AT- and GC-skewness as strong predictors to explore gene order variability within Octocorallia. While there is no evidence that ‘atypical’ AT/GC skewness values may lead to mitochondrial rearrangements, our data suggest that biases in GC/AT contents are often linked to mitogenome rearrangements in octocorals.

In this study we investigated the complete mitogenomes available for octocorals and updated the actual number of arrangements so far described to 12. Other studies have previously explored mitogenome diversity across the subclass, finding that even though different arrangements exist, all PCGs, ribosomal RNA and tRNA genes are conserved in four blocks (Brockman and McFadden 2012). Shimpi et al. (2017) proposed that synteny within these conserved blocks is likely due to the lack of recombination hotspots that promote genome rearrangements. However, our results indicated that rearrangements with non-conserved gene blocks are actually more than previously thought and the inversion and/or translocation of one or more of these blocks has led to seven different arrangements. Among the four types of rearrangements events (transpositions, inversions, inverse transpositions and tandem-duplication/random loss (TDRL)), inversions were the most frequent in the mitogenome evolution of octocorals. Although in nuclear genomes inversions are much more frequently observed than transpositions and inverse transpositions (Blanchette et al. 1999; Yancopoulos et al. 2005), TDRL seem to be the main mechanism driving rearrangement in mitogenomes, unlike inversions that, requiring recombination, are poorly documented in metazoan mitogenomes (Xia et al. 2016; Arndt and Smith 1998; Boore 2000; Osigus et al. 2013). Pante et al. (2013) and Hogan et al. (2019) reported, respectively, gene orders F (*Isidoides armata*) and G (*Umbellula* sp. 1) with non-conserved gene blocks. Similarly, we found five new gene orders (F1, H, I, J, K) characterised by non-conserved gene blocks. However, our findings, in agreement with what was previously discovered, showed that more than half of the existing mitogenome arrangements have non-conserved blocks. Additional studies on mitochondrial RNA processing and functional analyses involving, for example, the ability of organisms to perform recombination repair needs to be explored in order to better understand the reasons for multiple mitochondrial genome arrangements in octocorals.

Our data also showed the presence of gene order variability at the intrageneric level as *Phenganax stokvisi* has a different arrangement from congeneric *P. marumi* and *P. subtilis*. The occurrence of mitogenome rearrangements within single octocoral genera are uncommon, as they have only been detected among species of *Phenganax* and *Umbellula*, highlighting that new gene orders are generally conserved in a genus and/or a clade after the rearrangement event(s). However, in order to corroborate this finding further mitogenomes in species-rich genera need to be sequenced. According to our TreeRex analyses mitochondrial genome evolution in the subclass is discontinuous, with the rearranged gene orders restricted only to some regions of the phylogenetic tree. Such variability suggests, as supported by our ancestral state reconstruction, that different rearrangement events arose independently and that most likely the new gene orders, instead of being derived from other rearranged orders, came from the ancestral and most common gene order. This is further confirmed by the presence of at least one species with the ancestral gene order in each of the malacalcyonacean (excluding clade M1) and scleralcyonacean clades.

Regarding metazoan mitogenomes, Boore and Brown (1998) suggested that gene arrangements may imply common ancestry as the same gene order is unlikely to occur independently in separate lineages. We found that gene order variability among octocorals is not necessarily related to the taxonomy and phylogeny of the different taxa, as multiple arrangements have been found within the same family/genus, whereas the same arrangement can be in common across taxonomically and phylogenetically distinct genera, such as for instance the arrangement type ‘B’ which is shared by some sea pens and keratoisidids. As well, our mitogenomic investigations supported the interfamilial phylogeny in Keratoisididae where the S1 clade diverged earlier and retained the ancestral gene order arrangement. The recently published papers on mito-nuclear discordance within Anthozoa (Quattrini et al. 2023) and evidence for positive selection impacting the evolution of octocoral mitogenomes (Ramos et al. 2023) remark on the limits of using mt-PCGs or complete mitogenomes to infer accurate species-level phylogenies, yet here we observed that the study of mitochondrial gene orders is useful to explore the evolution of octocorals, and in some cases data can be exploited to address the taxonomy or to assess the phylogenetic placement of a given taxon. This was the case for *Protodendron*, whose gene order arrangement supported its phylogenetic placement within Xeniidae, which is otherwise difficult to predict based on its morphology. Complete mitogenome sequences are now available for one or more representatives of most families of octocorals. If additional arrangements exist, they will likely be restricted to particular genera or species. Further study of cases where multiple different arrangements have evolved within the same family or genus may help shed further light on the drivers and mechanisms of mitogenome evolution in non-bilaterian taxa.

## Acknowledgements

The first author is thankful to the National Institute of Genetics (NIG) for providing access to a supercomputer.

## Supplemental material

**Supplementary Table S1**

List of the specimens used for the analyses including taxonomic information, accession number, gene order, mitogenome size, GC% and AT-GC-skewness.

